## Supplementa for "Disentangling plant- and environment-mediated drivers of active rhizosphere bacterial community dynamics during short-term drought"

### Methods: Accounting for phantom taxa and designating the active community members

After generation of OTU and taxonomy tables, all subsequent downstream processing were performed in R version 4.0. The R package decontam [1] revealed the library sizes for the negative controls and the true samples and helped to determine the number and identity of contaminants in the dataset (**Figures S3A** and **S3B**). This package was used to remove contaminants from the dataset by using the prevalence method. Contaminating OTUs, mitochondria, and chloroplast sequences were filtered from the datasets. Based on rarefaction curves, a subsampling depth of 15,000 reads was selected for both datasets (**Figure S3C**). After subsampling to equal depth of 15,000 reads, 16S rRNA to rRNA gene ratios (hereafter, 16S rRNA:rRNA gene) were computed from the DNA and cDNA datasets based on two methods as described in [2]; specifically method 1 and method 2 therein. Briefly, these methods differed in their level of conservativeness when accounting for OTUs that have sequences detected in the cDNA (suggesting active transcription at the time of sampling), but no detected sequences in the DNA. Thus, these taxa were potentially in low abundance so that the DNA sequencing was not sensitive enough to detect them, but their transcripts were in relatively higher abundance and could be readily detected given the sequencing depth. Method 1 sets the ratio for all OTUs with cDNA>0 and DNA=0 to 100, and method 2 computes the ratio for all OTUs with cDNA > 0 and DNA= 0 by setting the DNA=1. Both methods produced similar results for all ecological statistics when active taxa were designated at a ratio threshold >=1. When ratio threshold was set to >1, method 1 overestimated the phantom OTUs remaining compared to method 2. However, we proceeded with ratio threshold >=1 so we could account for all those taxa which were present in equal abundances in DNA and cDNA datasets, and hence contributed to a ratio=1. Given that similar results were observed at this threshold for both methods, we picked method 2 to account for phantom taxa. Overall trends in beta diversity did not change, whether singletons were removed or kept, or whether the phantom taxa were included or excluded. In general, most of the phantom OTUs in the rarefied dataset (>10,000 out of 14,479 phantom taxa; total OTUs in rarefied dataset was 48,647) had very low occupancy across all the 212 samples in the cDNA dataset. We define phantom taxa here as those OTUs whose DNA counts (row sums) across samples is equal to zero, but cDNA counts (row sums) are greater than zero. For around 8000 phantom OTUs, around 100% of all instances of detection in the cDNA dataset across 212 samples had cDNA read equal to 1. Thus, for the purpose of analyzing the active community data, we kept the singletons and filtered to only those phantom taxa that were detected in at least 5% of the samples in the cDNA dataset. Around 922 taxa out of the 14,479 phantom taxa (6.4%) in the rarefied dataset were detected in the DNA reads of the unrarefied dataset, indicating that rarefaction could have led to the loss of DNA detected in the final dataset, though that percentage is very small. For those phantom taxa detected in the unrarefied dataset, the distribution of DNA counts revealed that most of those taxa were rare with reads summed across samples mostly adding up to less than 20 reads. Not surprisingly, only 0.04% of the total DNA reads in the unrarefied dataset were attributed to those 922 phantom taxa. The occupancy of most of those 922 phantom taxa in the unrarefied DNA dataset were also less than 1%. After selecting phantom taxa based on the threshold value of 5% occupancy across cDNA samples, 214 phantom OTUs remained out of the 14,479 taxa. A total of 12,388 OTUs remained after filtering by 16S rRNA:rRNA gene ratio threshold >=1 and selecting phantom taxa at 5% detection. The abundances of these OTUs were filtered to the DNA counts (for phantom taxa this was set to 1) and all analyses were performed based on this final active dataset filtered to DNA abundance. Consequently, while every sample was initially rarefied to 15,000 reads, each sample's active community varied slightly in total reads.

**Methods: Selection of active OTUs**

The percent phantom taxa in the rarefied dataset were assessed, i.e., those taxa which have RNA reads but lack DNA reads. Around 30-50 % of all OTUs were phantom taxa in the bean and switchgrass dataset. Roughly 10% of all OTUs were singleton phantom taxa in both switchgrass and bean dataset.

We analyzed the percent active OTUs in the dataset after using the two methods for ratio calculation; method 1: setting all cDNA>0 and DNA=0 to ratio 100 and method 2: setting all DNA= 0 to DNA=1. The percent active OTUs maintained the same overall trend across sampling days for both bean and switchgrass planted and unplanted treatments irrespective of method and ratio threshold (>1 versus >=1) with slight decreases in active OTUs when threshold was > 1 compared >=1 (associated code to reproduce results is included**).**

**Results: Metabolome analysis of switchgrass**

To unravel the global differences between the metabolomes of switchgrass rhizosphere and bulk soil before and after the drought treatment, we carried out untargeted LC-MS based metabolomics analysis of soils collected during the drought-treatment experiment. In addition, we also profiled the global changes occurred in metabolomes of switchgrass shoot and root caused by the drought treatment (**Figure S2**). The metabolite profiling revealed a total of 3,532 distinct metabolite features whose maximum abundance among the biological samples was ≥ 500 (**Data 1 - 3, Supplemental Data File**). A comparative analysis revealed 1051 root-, 538 shoot- and 231 soil- specific features (**Figure S4A**). Root and shoot shared 1551, root and soil shared 883 and leaf and soil shared 750 features, while 736 features were shared by all three groups (**Figure S4A**). The normalized abundance of features for soil were used for principal component analysis (PCA), which revealed the drought and watered planted (rhizosphere) soil separating from the rest soil types, including all the unplanted (bulk) soil and planted soil on the day 0 (the "Pre-drought" condition, **Figure S4B**). These metabolite differences indicate that the soil metabolites were altered by planting the switchgrass after five days. To further explore the changes occurring in the metabolome of drought-treated planted soil, a scatter plot based on PCA scores was used. The result showed some divergence between the drought and watered planted soil samples (**Figure S4C**), but this was not statistically supported. A variable of importance (VIP) coefficient was generated from the PLS-DA model for each metabolite feature. The top 15 most differentially accumulated features (with the highest VIP coefficient) between the different types of soil are shown in **Figure 3**. The feature, *10.11_1000.5271n*, has the highest VIP coefficient indicating it as the most important variable among altered metabolites for separation of the metabolomes (**Figure 3**). The LC-MS/MS spectra revealed this metabolite feature is a previously identified switchgrass mono-glycosylated steroidal saponin [3] (**Figure S5)**. When the top 50 features with the highest VIP coefficients were used to generate a heatmap with hierarchical clustering, the planted and unplanted soil samples formed two distinct clusters that separated from the other soil samples including the pre-drought planted and unplanted soil (**Figure S4D**). Among these important features, a few of them were annotated as the specialized metabolites, including a few previously reported switchgrass saponins [3] and root accumulating diterpenoids with elevated concentrations in the drought treated planted soil samples [3, 4] (**Data 4, Supplemental Data File**). The higher abundances for these specialized metabolites in the drought planted soil implied that switchgrass might release them into the rhizosphere soil carrying out certain biological functions when stressed by drought.

**Table S1.** Chemical and physical analysis of field-collected soils for bean and switchgrass.

| Soil Sample | Switchgrass (*Panicum virgatum*) | Bean (*Phaseolus vulgaris*) |
|---|---|---|
| Soil pH | 6 | 6.7 |
| Lime Index | 69 | 71 |
| Phosphorus (ppm) | 21 | 200 |
| Potassium (ppm) | 72 | 109 |
| Magnesium (ppm) | 124 | 88 |
| Calcium (ppm) | 913 | 574 |
| CEC (meq/100g) | 7 | 3.9 |
| K | 3.2 | 7.2 |
| Mg | 17.9 | 18.9 |
| Ca | 78.9 | 73.9 |
| Organic Matter (%) | 2 | 1.1 |
| Nitrate-N (ppm) | 3.5 | 2.5 |
| Ammonium-N (ppm) | 1.6 | 0.6 |
| Soil moisture (%) | 14.1 | 13.7 |

**Table S2.** Water addition schedule by treatment for switchgrass.

| Treatment date (Year-2019) | Water added for watered treatment | Water added for drought treatment | Sampling day (sampling done without watering that morning) |
|---|---|---|---|
| Sat Apr 27 | 10 ml | 5 ml | Pre-drought – Day 0 |
| Sun Apr 28 | 0 ml | 0 ml | |
| Mon Apr 29 | 10 ml | 3 ml | Day 2 |
| Tues Apr 30 | 0 ml | 0 ml | Day 3 |
| Wed May 1 | 10 ml | 1 ml | Day 4 |
| Thurs May 2 | 10 ml | 0 ml | Day 5 |
| Fri May 3 | n/a | n/a | Day 6 |

**Table S3.** Water addition schedule by treatment for bean.

| Treatment Date (Year-2019) | Water added for watered treatments | Water added for drought treatments | Sampling day (sampling done without watering that morning) |
|---|---|---|---|
| Wed, May 22 | 15ml morning, 15ml afternoon. | 8ml morning, 8ml afternoon | Pre-drought – Day 0 |
| Thurs, May 23 | 15ml morning, 15ml afternoon. | 6ml morning, 6ml afternoon | |
| Fri, May 24 | 15ml morning, 15ml afternoon. | 4ml morning, 4ml afternoon | Day 2 |
| Sat, May 25 | 15ml morning, 15ml afternoon. | 2ml morning, 2ml afternoon | Day 3 |
| Sun, May 26 | 15ml morning, 15ml afternoon. | 0ml morning, 0ml afternoon | Day 4 |
| Mon, May 27 | 15ml morning, 15ml afternoon. | 0ml morning, 0ml afternoon | Day 5 |
| Tues, May 28 | n/a | n/a | Day 6 |

**Table S4.** Differences in gravimetric soil moisture and shoot biomass by experimental factor as assessed by a three-way ANOVA with interactions (Type III). The F-statistics are reported with significance indicated at p<0.05*, p<0.01**, and p<0.001***.

| Factor | Gravimetric soil moisture | | Shoot biomass | |
|---|---|---|---|---|
| | Bean (n=100, without pre-drought samples) | Switchgrass (n=100, without pre-drought samples) | Bean (n=50, without pre-drought samples) | Switchgrass (n=50, without pre-drought samples) |
| planted | 1160.21 *** | 6.66 * | | |
| drought | 165.49 *** | 28.43 *** | 21.18 *** | 0.30 |
| Sampling day | 7.52 *** | 0.20 | 1.89 | 3.43 * |
| planted:drought | 31.46 *** | 0.00 | | |
| planted:sampling day | 3.41 * | 1.01 | | |
| drought:sampling day | 11.97 *** | 0.24 | 1.49 | 1.32 |
| planted:drought:sampling day | 2.84 * | 0.39 | | |

**Table S5.** Permuted analysis of variance analysis to determine differences in microbiome structure given experimental treatments. Sums of squares, degrees of freedom, R-squared values and Pseudo F values are reported. The significance of each test is indicated at p<0.05*, p<0.01**, and p<0.001***.

| Factor | Switchgrass (n=97, without pre-drought samples) | | | | Bean (n=96, without pre-drought samples) | | | |
|---|---|---|---|---|---|---|---|---|
| | Sums of squares | Degrees of freedom (DF) | $R^2$ | Pseudo-F | Sums of squares | Degrees of freedom (DF) | $R^2$ | Pseudo-F |
| planted | 0.56 | 1 | 0.03 | 3.30*** | 3.12 | 1 | 0.17 | 21.57*** |
| drought | 0.28 | 1 | 0.02 | 1.64*** | 0.39 | 1 | 0.02 | 2.71*** |
| sampling day | 0.87 | 4 | 0.05 | 1.28*** | 0.95 | 4 | 0.05 | 1.65*** |
| planted:drought | 0.19 | 1 | 0.01 | 1.13 | 0.27 | 1 | 0.01 | 1.84* |
| planted:sampling day | 0.79 | 4 | 0.05 | 1.16* | 0.75 | 4 | 0.04 | 1.29* |
| drought:sampling day | 0.87 | 4 | 0.05 | 1.27** | 0.78 | 4 | 0.04 | 1.35* |
| planted:drought:sampling day | 0.74 | 4 | 0.04 | 1.08 | 0.68 | 4 | 0.04 | 1.17 |

**Table S6.** Differences in the number of observed active OTUs (richness) across experimental factors, as assessed using a three-way ANOVA with interactions (Type III tests). F values are reported and significance is indicated at p<0.05*, p<0.01**, and p<0.001***.

| Factor | Observed OTUs | |
|---|---|---|
| | Bean (n=96) | Switchgrass (n=97) |
| planted | 60.12 *** | 0.35 |
| drought | 0.07 | 1.99 |
| sampling day | 1.61 | 0.79 |
| planted:drought | 2.32 | 4.30 * |
| planted:sampling day | 2.12 | 0.49 |
| drought:sampling day | 1.23 | 1.16 |
| planted:drought:sampling day | 1.42 | 0.49 |

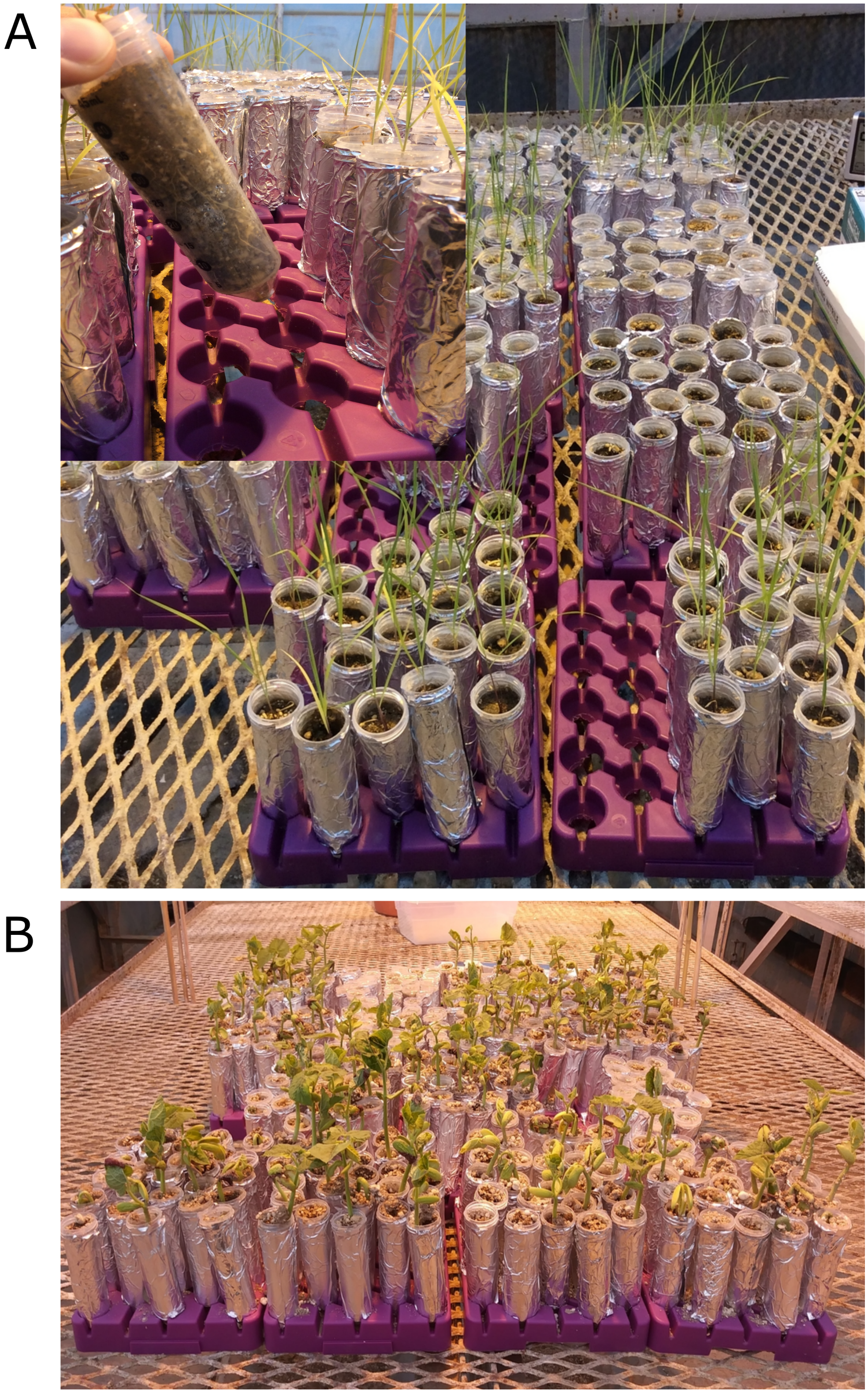

**Figure S1.** Photo of experimental set up in the greenhouse for A) switchgrass (inset- tube showing switchgrass roots grown to fill to the inside edge of the tube) and B) bean.

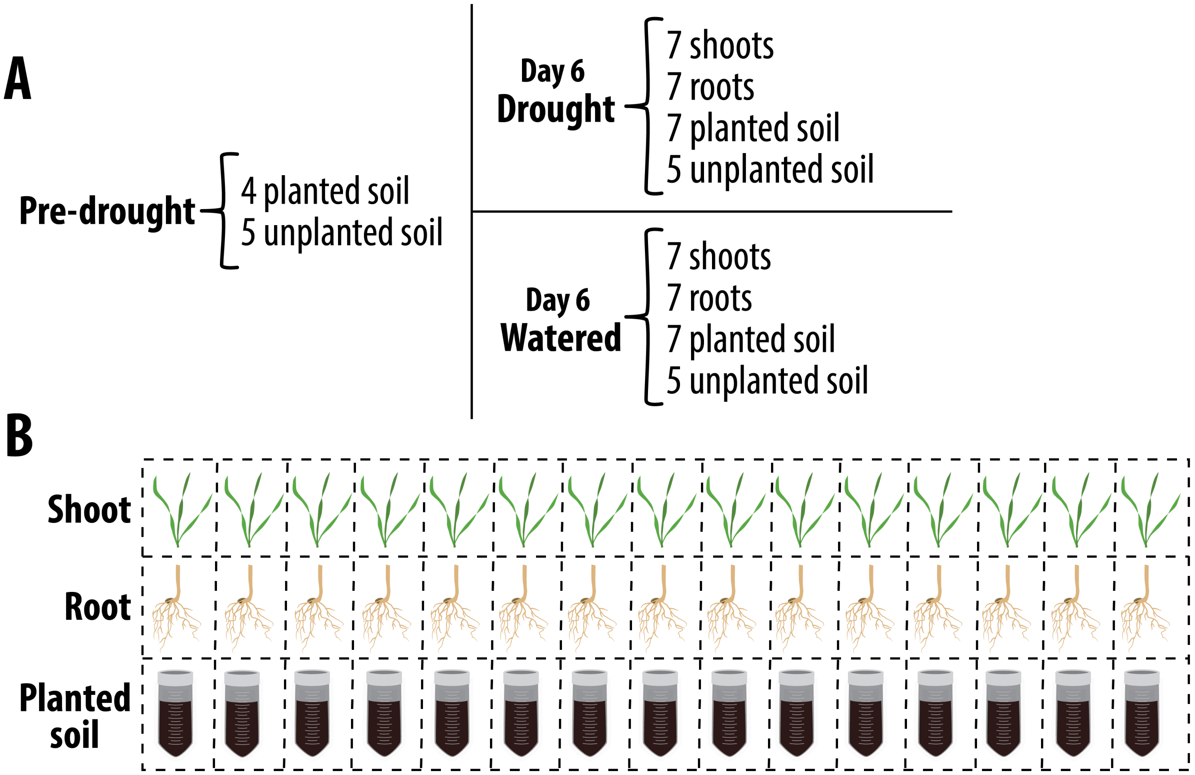


**Figure S2.** Sample collection for the switchgrass untargeted metabolomics. **A.** Sample panel with the numbers of replicates indicated. **B.** The drought and watered switchgrass tissue and planted soil replicates were combined from the samples that were collected from individual plants.

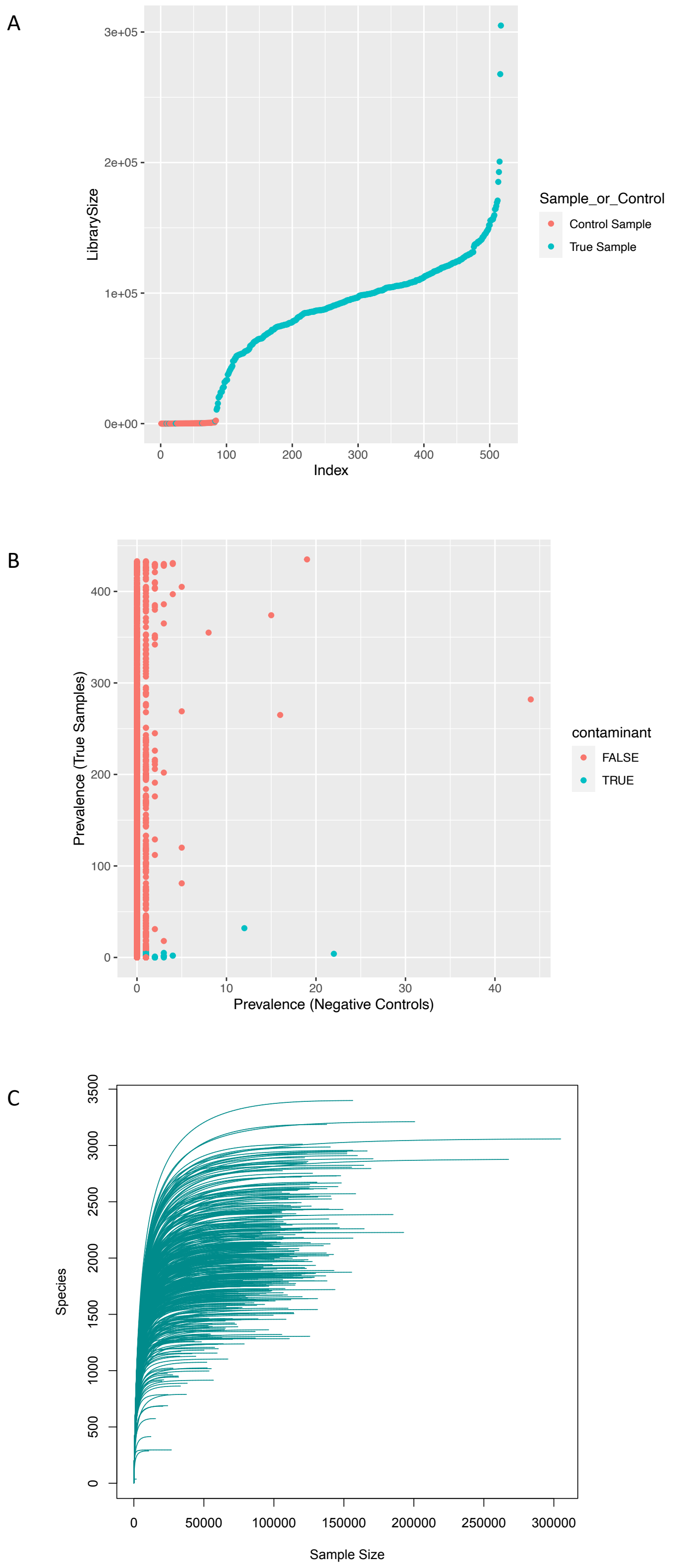
A
3e+05
2e+05
1e+05
0e+00
LibrarySize
0
100
200
300
400
500
Index
Sample_or_Control
Control Sample
True Sample
B
400
300
200
100
0
Prevalence (True Samples)
0
10
20
30
40
Prevalence (Negative Controls)
contaminant
FALSE
TRUE
C
3500
3000
2500
2000
1500
1000
500
0
Species
0
50000
100000
150000
200000
250000
300000
Sample Size

**Figure S3.** Sequencing quality control checks. (A) Library sizes of true samples (blue) and negative control (red) samples for both DNA and cDNA-sequenced samples. (B) OTUs designated as either true or false contaminants based on their prevalence in true samples and negative controls using package decontam in R. (C) Rarefaction curves for all DNA and cDNA samples (pre-rarefaction). Curves start to plateau out at 13-15k reads (n=438).

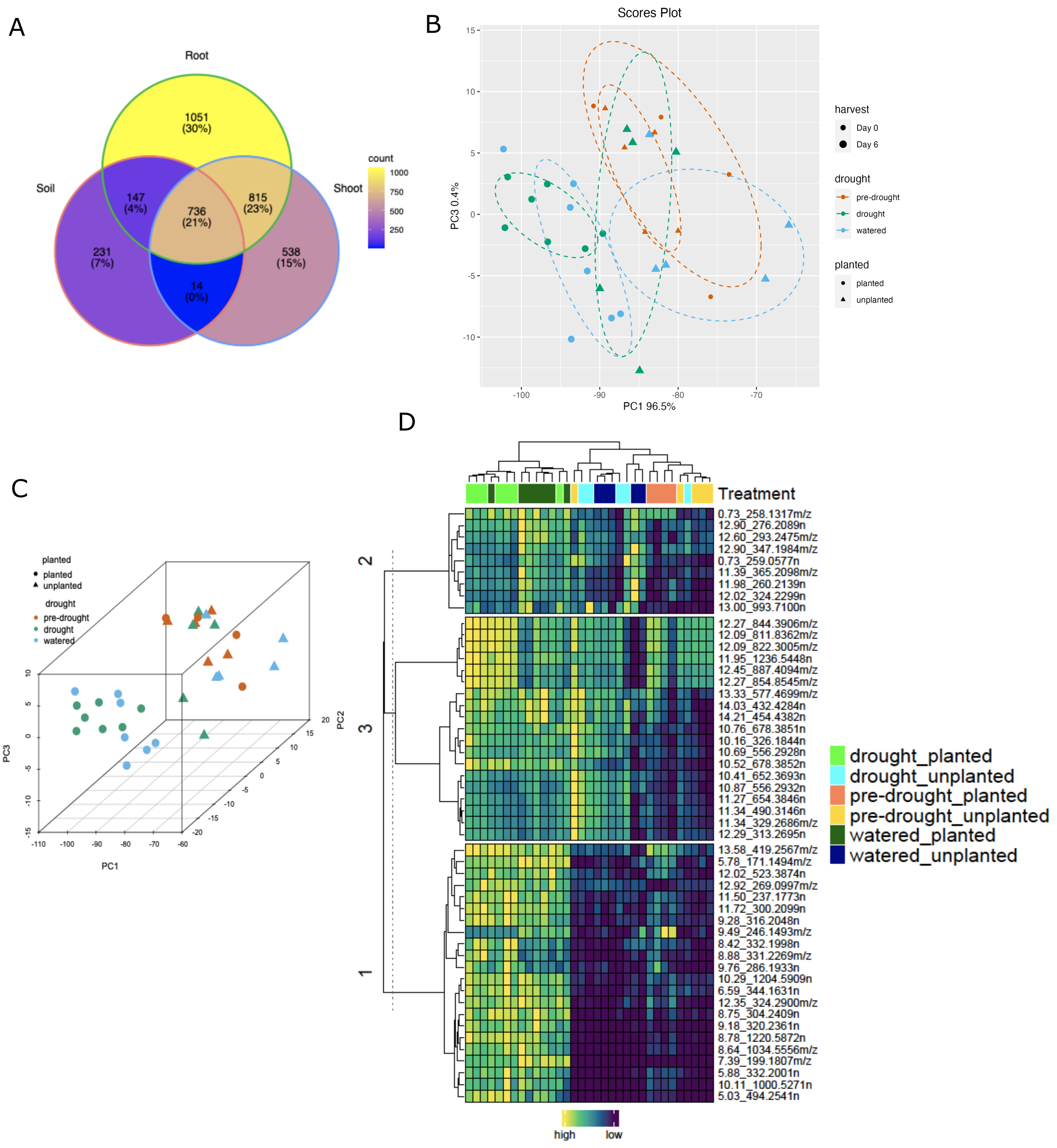


**Figure S4.** The untargeted metabolomics revealed metabolite differences between the soil types. A. Venn diagram documenting comparative metabolite profiles as well as number of specific and shared features identified in switchgrass root, shoot and soil. B. PCA scores plot of the metabolomes of different soil types showing the principal components PC1 and PC3. C. 3-D scatter plot of PCA scores plot of the metabolomes of different soil types showing PC1, 2 and 3. D. Heatmap showing relative abundances of the top 50 PLS-DA most important features (vertical axis) across the biological samples (horizontal axis).

For the hierarchical clustering of features, Euclidean and Ward.D were used as the distance measure and clustering method respectively (sample size root (n)=30, shoot (n)=30, soil(n)=49).

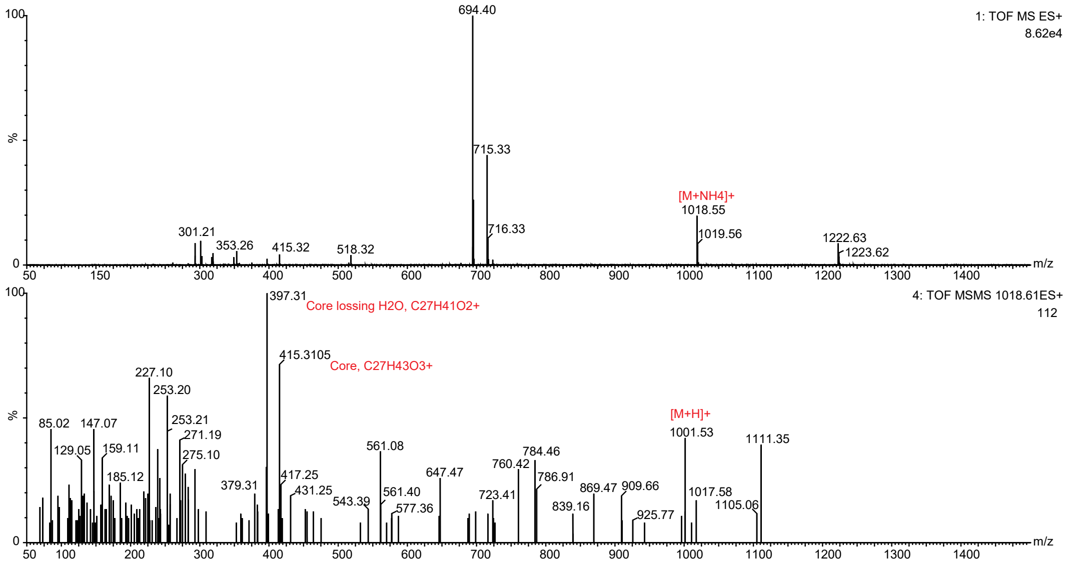


**Figure S5.** MS/MS spectra for the feature, *10.11_1000.5271n,* obtained by the positive LC-MS analysis, DDA mode. Top trace: survey scan; bottom trace: MS/MS. The molecular ions and sapogenin core fragment ions were annotated. M, molecular ion.

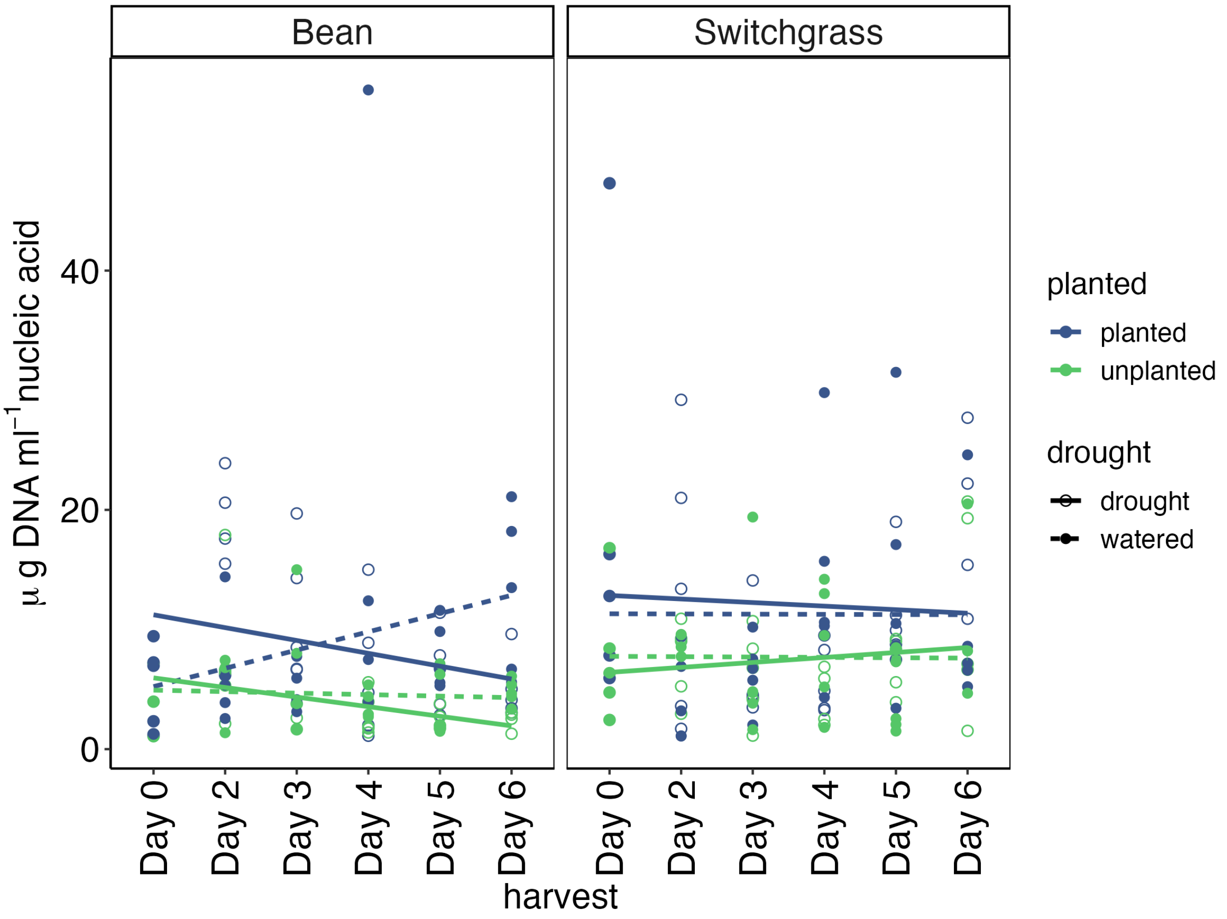


**Figure S6.** Recovered DNA concentrations of soil samples by crop, treatment, and time point. DNA was measured using Qubit 2.0 with the dsDNA BR assay kit.

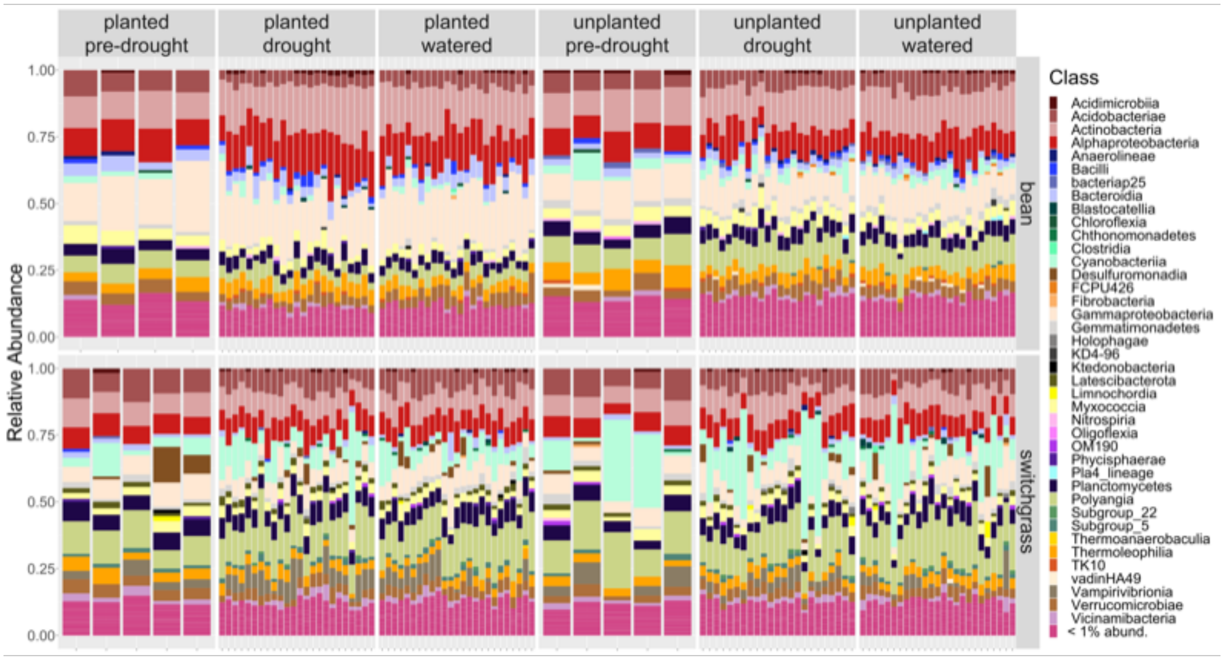


**Figure S7.** Stacked bar plots depicting the Class-level, active bacterial community composition by treatment for bean and switchgrass. All samples collected are shown (n=212).

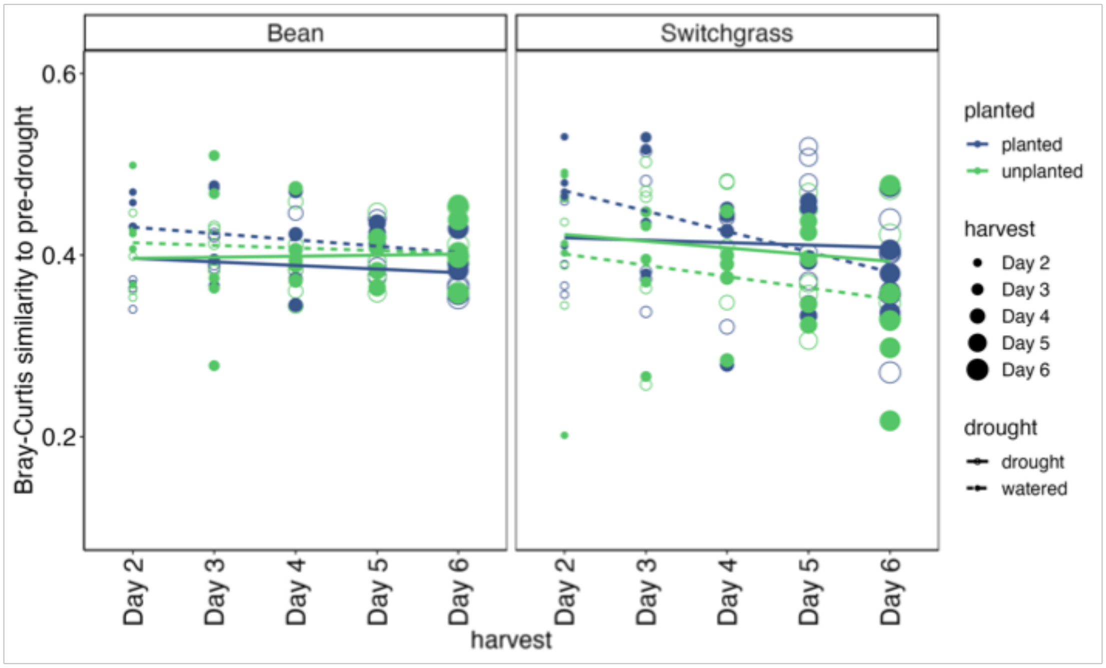


**Figure S8.** For each experimental treatment, changes in Bray Curtis similarity to the pre-drought community over time and/or with increasing severity. Sampling day is indicated by the symbol size, and treatment by colors (blue is planted, green is unplanted) and line type (solid is drought and dashed is watered). The linear regressions were not significant for any treatment, except for the watered, planted switchgrass (blue, dashed line on the right panel).

### **RE**FERENCES


1. Davis NM, Proctor DM, Holmes SP, Relman DA, Callahan BJ: Simple statistical identification and removal of contaminant sequences in marker-gene and metagenomics data. *Microbiome* 2018, 6(1):226.
2. Bowsher AW, Kearns PJ, Shade A: 16S rRNA/rRNA Gene Ratios and Cell Activity Staining Reveal Consistent Patterns of Microbial Activity in Plant-Associated Soil. *mSystems* 2019, 4(2).
3. Li X, Sarma SJ, Sumner LW, Jones AD, Last RL: Switchgrass Metabolomics Reveals Striking Genotypic and Developmental Differences in Specialized Metabolic Phenotypes. *Journal of Agricultural and Food Chemistry* 2022, 70(26):8010-8023.
4. Tiedge K, Li X, Merrill AT, Davisson D, Chen Y, Yu P, Tantillo DJ, Last RL, Zerbe P: Comparative transcriptomics and metabolomics reveal specialized metabolite drought stress responses in switchgrass (Panicum virgatum). *New Phytologist* 2022, 236(4):1393-1408.
